# The timing of hemodynamic changes reliably reflects spiking activity

**DOI:** 10.1101/269696

**Authors:** Ali Danish Zaidi, Niels Birbaumer, Eberhard Fetz, Nikos Logothetis, Ranganatha Sitaram

## Abstract

Functional neuroimaging is a powerful non-invasive tool for studying brain function, using changes in blood-oxygenation as a proxy for underlying neuronal activity. The neuroimaging signal correlates with both spiking, and various bands of the local field potential (LFP), making the inability to discriminate between them a serious limitation for interpreting hemodynamic changes. Here, we record activity from the striate cortex in two anesthetized monkeys (*Macaca mulatta*), using simultaneous functional near-infrared spectroscopy (fNIRS) and intra-cortical electrophysiology. We find that low-frequency LFPs correlate with hemodynamic signal’s peak amplitude, whereas spiking correlates with its peak-time and initial-dip. We also find spiking to be more spatially localized than low-frequency LFPs. Our results suggest that differences in spread of spiking and low-frequency LFPs across cortical surface influence different parameters of the hemodynamic response. Together, these results demonstrate that the hemodynamic response-amplitude is a poor correlate of spiking activity. Instead, we demonstrate that the timing of the initial-dip and the hemodynamic response are much more reliable correlates of spiking, reflecting bursts in spike-rate and total spike-counts respectively.

## Introduction

Functional neuroimaging is currently one of the best non-invasive tools for understanding brain function in healthy, and dysfunction in diseased subjects (Logothetis, 2008), where the peak amplitude of the hemodynamic response is most often used as a proxy for the strength of underlying excitatory neural activity, such as spiking or activity in the gamma band. However, the precise interpretation of the neuroimaging signal is rendered difficult by the fact that it has been reported to reflect various neuronal processes, such as multi-unit spiking, the activity in the various frequency bands of the LFP (Bartolo et al., 2011; Boynton, 2011; Goense and Logothetis, 2008; Logothetis et al., 2001; Maier et al., 2008; Murayama et al., 2010; Rauch et al., 2008; Viswanathan and Freeman, 2007), and even the relationships among different LFP bands (Magri et al., 2012). Spikes and low-frequency LFPs may, however, represent distinct neural processes such as feedforward and feedback influences (Bastos et al., 2015). For example, spiking seems to represent the features of a visual stimulus (feedforward), whereas neuro-modulatory inputs are reflected in low-frequency LFP bands (feedback) (Belitski et al., 2008). This makes the inability to differentiate between activations associated with spiking-activity or low-frequency LFPs the major limitation in the interpretation of hemodynamic changes (Logothetis, 2008). However, the major portion of such studies focus mainly on the amplitude of the hemodynamic signal, and two other parameters, namely, its peak-time (the time taken by the hemodynamic response to reach its peak) and initial-dip (a transient dip in the hemodynamic signal observed just after stimulus onset) have mostly been ignored.

Interestingly, spiking and low-frequency LFPs are known have different spatio-temporal features. For example, low-frequency LFP activity is more temporally synchronized, but has a larger spread over cortical surface (Logothetis et al., 2007). Spiking activity, in contrast, is spatially more localized, but has a larger temporal spread (e.g. a train of spikes). Since the hemodynamic response is believed to reflect the spatio-temporal pattern of local neuronal activity, it isn’t unreasonable to assume that such differences in the pattern of neuronal activity might be reflected in different features of the hemodynamic signal. The ability to identify features in the hemodynamic signal that serve as markers for a specific kind of neuronal activity would enable a clearer and much more precise interpretation of functional neuroimaging studies. Hence, to determine how different hemodynamic parameters correlated to different neuronal processes (such as spiking or LFPs), we performed simultaneous measurements of hemodynamic signals (using fNIRS) and intra-cortical electrophysiology in the primary visual cortex (V1) of two anesthetized monkeys (**Fig. 1a-b**). FNIRS uses a light emitter-detector pair (optode pair) to measure changes in the concentration of oxygenated ([HbO]) and deoxygenated ([HbR]) hemoglobin in a small volume of tissue (Ferrari and Quaresima, 2012; Villringer, 1997). We have recently demonstrated that fNIRS has high SNR when acquired epidurally (Zaidi et al., 2015), making it ideal for studying local neuro-vascular interactions. Using this technique, we recorded sessions of both spontaneous and stimulus-induced activity in the primary visual cortex of two anesthetized monkeys, where a high-contrast whole-field rotating chequerboard was used as visual stimulus. **Fig 1c** shows traces of [HbO], [HbR] and spiking activity for an example run consisting of 20 trials. Each trial consisted of 5s of stimulus presentation followed by 15s of a dark screen. The visual stimulus invoked strong responses, clearly observed in single trials as changes in the traces of HbO, HbR and spike rates.

**Figure 1.**
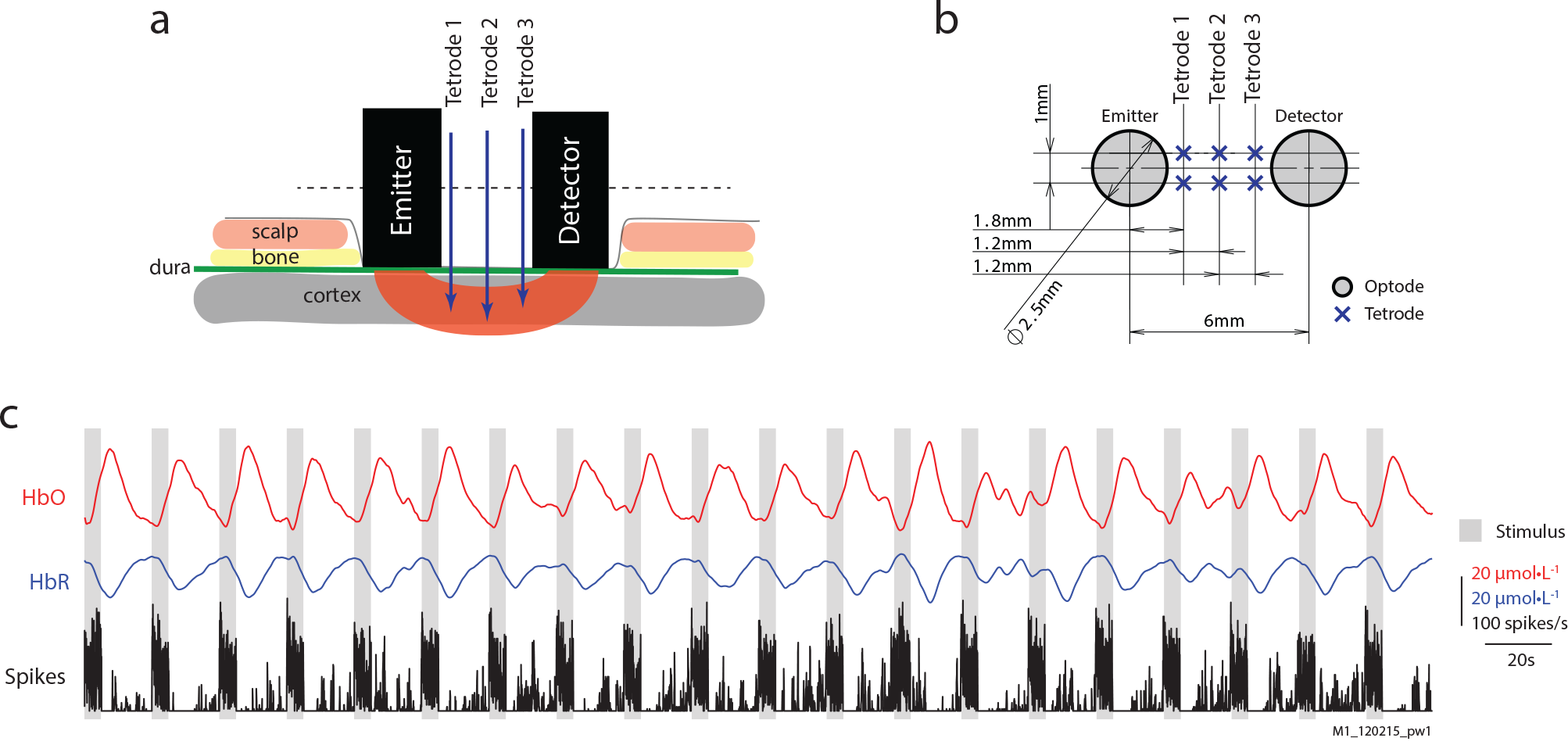
Overview of acquired signals. A) Illustration of the sensor array with placement of fNIRS optodes and electrodes relative to scalp and brain tissue. B) Transverse section of the sensor array with distances between optodes and electrodes. See methods for details. C) Traces of HbO, HbR, and Spiking from an example run with 20 trials. Grey bars represent epochs of visual stimulation.

## Results

### Timing of the hemodynamic changes best reflects spiking activity

To analyze the relationship between neural modulation and the hemodynamic response, we sorted the trials based on the strength of their spike-rate modulation (**Fig. 2a_1-2_**), and obtained their corresponding HbO traces (**Fig. 2a_3_**). A negative trend was observed in the peak-amplitude of the HbO response (**Fig 2a_3_**), whereas a positive trend was observed in the HbO peak-time (**Fig 2a_4_**). To overcome the large trial-by-trial variation in the data, we sorted the trials from low to high spike-rate modulation and binned them into five groups. The mean spike rates and [HbO] traces for each group are shown in **Fig. 2b_1-2_**. **Fig 2b_3-4_** show the means spike rate versus the mean HbO peak-amplitude and HbO peak-time for each group, along with their standard error of mean (SEM). Surprisingly, spike-rate modulation correlated negatively with HbO peak-amplitude (**Fig. 2b_3_**; r =−0.98; p< 0.005; n = 50/group, 5 groups), but positively with HbO peak-time (**Fig. b_4_**; r = 0.93, p < 0.023; n=50/group, 5 groups). This relationship was also observed for [HbR] signals (albeit weaker) and was independent of both the number of groups the data were divided into (*supplementary figure 1*), and the threshold for spike detection (*supplementary figure 2*). Trials with higher spike-rate modulations also elicited higher peak-rates and total spike-counts during stimulus presentation (**Fig. 2b_1_**). A strong relationship between peak spike-rate during stimulus onset and HbO peak-time (r = 0.8; p < 0.01; **Fig. 2c_1_**; n=25/group, 10 group) was also observed. However, correlations between total spike-count during stimulus presentation and the HbO peak-time were the strongest (**Fig. 2c_3_**; r = 0.9; p < 0.001; n = 25/group, 10 groups).

**Fig 2.**
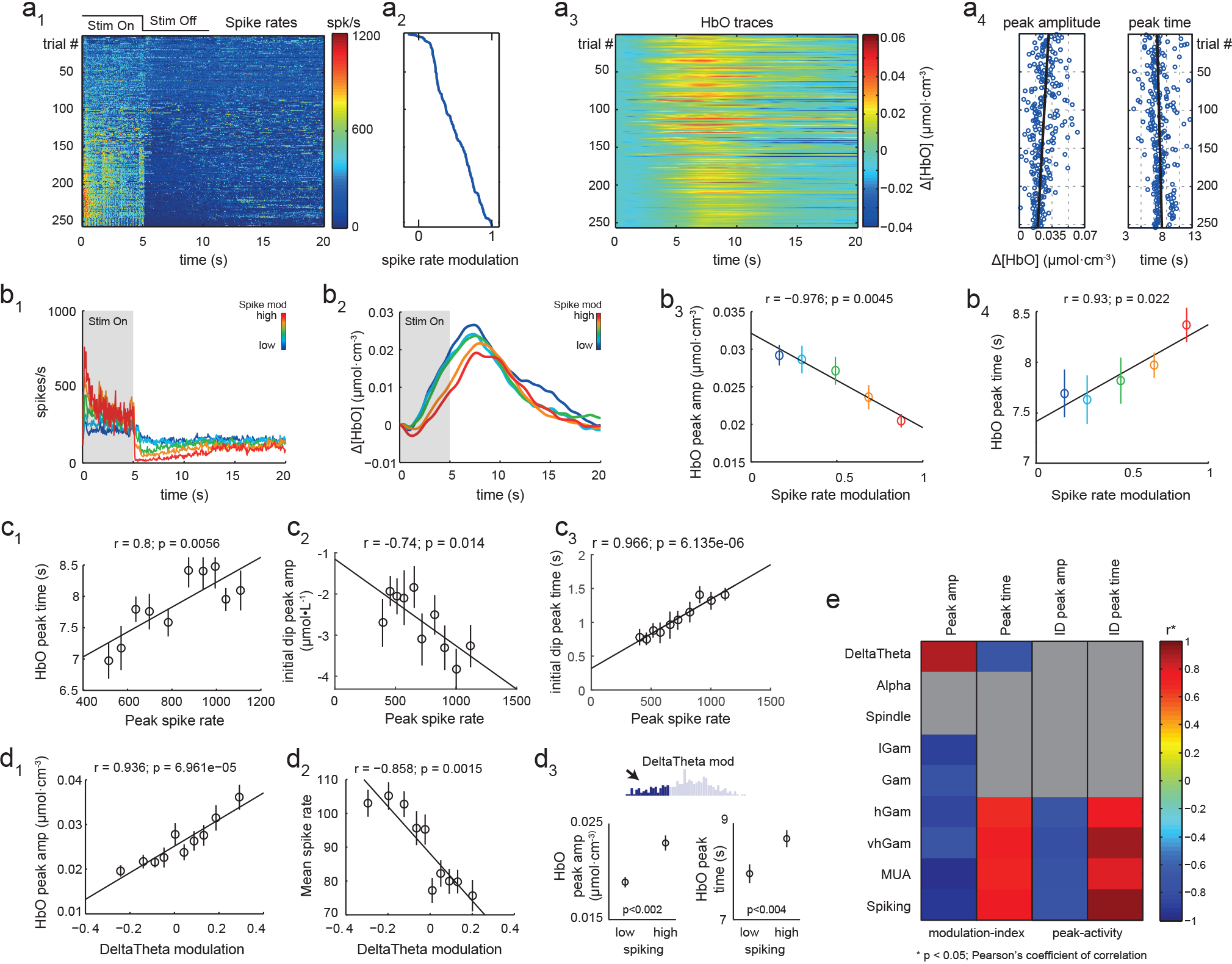
Correlation of spike-rates and LFPs to peak-time and peak-amplitude. a1) Spike rate of trials sorted based on modulation index of spike rate. a2) modulation index of each trial corresponding to a1. a3) [HbO] traces of the corresponding trials from a1. a4) A clear negative trend in peak-amplitude and positive trend in HbO peak-time can be observed for traces corresponding to a2, along with a large trial-by-trial variation. b1-2) Mean traces of spike rates (b1) and [HbO] traces (b2) for each group, when trials were sorted and grouped based on spike-rate modulation. b3-4) Correlation of mean spike-rate modulation with mean HbO peak-amplitude (b3) and mean peak-time (b4) (n=50 per group) from each group. c1-2) Correlations of mean peak-time with peak spike rate. c2-3) Correlations of peak spike-rate with initial-dip peak-amplitude and peak-time. d1) Correlation of DeltaTheta modulation with peak-amplitude (d1) and mean spike rate (d2) for trials sorted and grouped on DeltaTheta modulation. d3) Within the first quartile of DeltaTheta modulations (inset), spiking significantly affected both peak-amplitude and peak-time (p-values based on two-tailed Wilcoxon ranked sum test). e) Correlations of HbO response peak-amplitude and peak-amplitude with the modulation index, and the initial-dip peak-amplitude and peak-time with peak activity in different LFP frequency bands. Only significant correlations (p<0.05; n=25/group, 10 groups) are coloured. All error bars represent SEM.

We also observed robust initial-dips in the high spiking trials (**Fig. 2b_2_**). The initial-dip has been shown to represent local neuronal activity better than the positive peak (Kim et al., 2000; Watanabe et al., 2013). We therefore analyzed the relationship between spiking and both the initial-dip peak-amplitude and peak-time. We observed a strong correlation between the peak spike-rate and both the HbO initial-dip amplitude (r = −0.741; p = 0.014; n = 25/group, 10 groups; initial-dip amplitude is negative) and the initial-dip peak-time (**Fig. 2c_2_**; r = −0.966; p < 10^−5^; n=25/bin, 10 bins), demonstrating that although both the amplitude and timing of the initial-dip reflect bursts in spiking activity, its peak-time has a much stronger relationship. Among the various LFP bands, only the high frequency bands correlated with the peak-time of the initial-dip, but not with its amplitude (p<0.1). These results imply that for both the initial-dip and hemodynamic response, their peak-times reflect high-frequency neuronal activity more reliably than peak-amplitude, with the strongest correlations observed with spiking activity.

### Amplitude of the hemodynamic response reflects low-frequency LFP activity

We next determined the relationship between modulations in the various LFP bands and hemodynamic responses. We filtered the extra-cellular field potential into eight frequency bands, and obtained their respective band-envelope. The trials were then sorted and binned based on the strength of modulation in each band. This band modulation was then correlated with HbO peak-amplitude and HbO peak-time. A positive correlation between DeltaTheta (1-8 Hz) modulation and HbO peak-amplitude was found (r = 0.9, p < 0.001; **Fig. 2d_1_**; n=25/group, 10 groups; *supplementary figure 3*), and a negative (albeit non-significant) correlation with HbO peak-time (r = −0.5, p > 0.1; n=25/group, 10 groups). Opposing relationships of HbO peak-amplitude with spike-rates and DeltaTheta suggest an interaction between these two neuronal processes. Indeed, a strong negative correlation between DeltaTheta modulation and spike rates was observed (r = −0.86; p < 0.0015; **Fig. 2d_3_**). However, when comparing low and high spiking trials within the first quartile of DeltaTheta modulations, we observed a significant increase in both HbO peak-amplitude (p<0.002, n=62; Wilcoxon’s rank sum test) and HbO peak-time (p<0.004, n=62; Wilcoxon’s rank sum test) with increase in spiking activity (**Fig. 2d_3_**). This demonstrates that although spiking does affect HbO peak-amplitude positively, this effect is overshadowed by the influence of low frequency processes on both HbO peak-amplitude and spiking. **Fig. 2e** summarizes the relationship between modulations in different LFP frequency bands and the hemodynamic response parameters. Only modulations in DeltaTheta and Alpha (r = 0.69, p < 0.029; n = 25/group, 10 groups) correlated significantly with HbO peak-amplitude (and among themselves, r = 0.91, p < 0.001; n = 25/group, 10 groups). In contrast, only modulations in the MUA band correlated significantly with HbO peak-time. Since the modulation index of the DeltaTheta envelope was not significantly different than zero in our data (p > 0.1, Wilcoxon’s signed rank test, mean = 0.0149 ± 0.011 SEM; *supplementary figure 4*), these observations seem to reinforce the notion that spiking and low-frequency LFPs might represent different neural processes in the primary visual cortex (Belitski et al., 2008).

### Analysis of spontaneous activity reveals similar relationships

It might be argued that these relationships do not represent local neurovascular coupling, but are artifacts arising from strong visual stimulation. To ensure this was not the case, we analyzed sessions of spontaneous activity, recorded in the absence of visual stimulation with the subject’s eyes closed. Bursts in spike rates were detected and corresponding [HbO] traces were obtained from −5 to +20 seconds relative to the burst peak, and zero corrected by subtracting the mean value from −5 to −3s relative to burst peak. These traces were then sorted and grouped based on burst strength (Fig 3a1-2). The peak-rate of spiking was strongly correlated with both HbO peak-time (r = 0.97, p < 10−5; n=85/group, 10 groups; Fig 3b1) and initial-dip (r = −0.8, p = 0.005; n=85/group, 10 groups; Fig 3b2), but not HbO peak-amplitude (r = 0.31 p = 0.62 n = 85/group, 10 groups). The correlation between peaks in the different LFP band-envelopes and HbO peak-amplitude and HbO peak-time was also analyzed. Strong correlations were observed only between activity in high-frequency LFPs and HbO peak-time, whereas no significant relationships were found for peak amplitudes (Fig. 3c). These observations demonstrate that the relationship between spiking activity and hemodynamic peak-time are consistent across recordings of both spontaneous and stimulus-induced activity.

**Fig 3.**
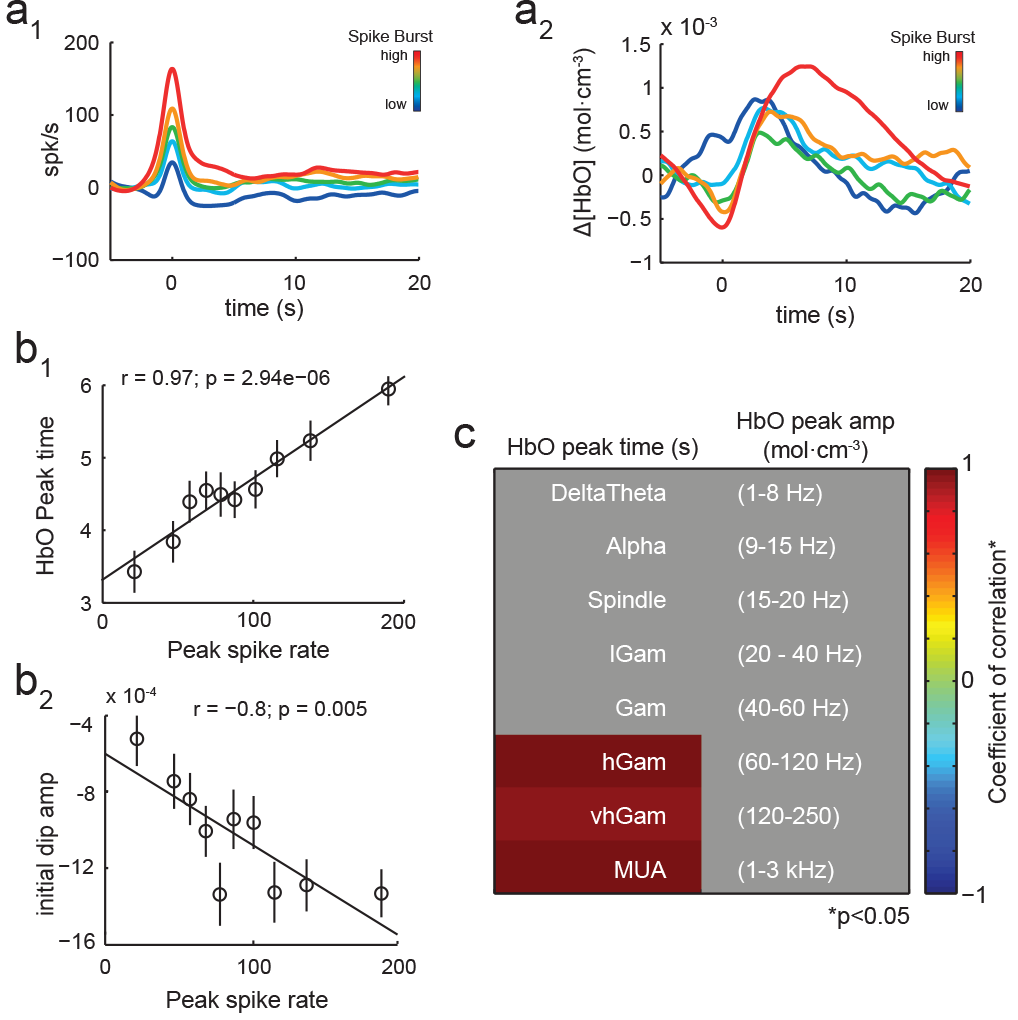
Relationships between neural activity and HbO parameters in spontaneous activity. a1-2) Mean traces of spike rate (a1) and corresponding [HbO] traces (a2) for each group, sorted from low to high spike-burst strengths. b) Correlations of the peak spike-rate for each group and the peak-time (b1) and initial-dip (b2) when spike bursts were sorted on peak spike rate. Error bars represent SEM. c) Correlation of peak-amplitude and peak-time with peaks in the envelope of different LFP frequency bands. Only significant correlations (p<0.05) are coloured. The peak-amplitude correlates with peaks in the Alpha band, and peak-time with peaks in the high gamma bands and MUA band.

## Discussion

*Prima facie,* our results might seem different from earlier studies of neurovascular coupling (Goense and Logothetis, 2008; Logothetis et al., 2001). However, it must be noted that most previous studies have analyzed time-series cross-correlations between neuronal activity and hemodynamic signals to estimate the strength of neurovascular coupling. In our study, we use a feature based approach, where individual features of neuronal activity are correlated with features of the hemodynamic response. Indeed, a cross-correlation based analysis reveals results similar to those previously observed (Goense and Logothetis, 2008; Logothetis et al., 2001), with low-frequency processes eliciting weak, and high-frequency processes eliciting strong correlations, for both time-series cross-correlation analysis (*Supplementary figure 5*) and HRF-convolutions obtained from spontaneous activity (*Supplementary figure 6*). However, cross-correlations are strongly dependent on the shape of the underlying signals, and consequently, may misrepresent underlying relationships. Our feature-based approach enables a more robust investigation, as we use multiple features of neuronal activity (e.g. neuronal modulation index, peak neuronal activity, mean activity during stimulus presentation, etc.) and obtain similar results.

These results probably arise from the differences in the spatio-temporal spread of low and high-frequency neuronal processes across the cortical surface. Both previous work (Logothetis et al., 2007) and our data demonstrate that correlations in high-frequency neuronal processes have a small spread over cortical surface (**Fig. 4a**) than those in high-frequencies (**Fig. 4b**). Modulations in lower frequencies, in contrast, are more synchronized (small temporal spread) and have a larger spatial-spread over the cortical surface, potentially dilating more vessels simultaneously. The simultaneous increase in flow rate of HbO would translate to an increase in the peak-amplitude of the hemodynamic signal. Indeed, both previous work (Devor et al., 2005) and our data (**Fig. 4c**) demonstrate that a larger spread of neuronal activity across the cortical surface causes larger hemodynamic response amplitudes. In contrast, localized neuronal responses are known to cause arteriole dilation with high spatial precision and limited spread (Sirotin et al., 2009). Being more localized, high-frequency processes such as spiking would dilate fewer arterioles, and with longer activity, the flow rate in these dilated arterioles would saturate. Thus, larger spike counts should affect HbO peak-time more than HbO peak-amplitude. Indeed, compared to early-peaking trials, late-peaking trials had significantly higher spike-rates during both the ‘On’ and ‘Off’ epochs (**Fig. 4d**). Furthermore, we found a strong correlation between the HbO peak-time and the total number of spikes up to the HbO peak-time for each trial (r=0.98, p<10^−6^, **Fig. 4d** **inset**). This result clearly demonstrates the HbO peak-time reliably reflects the sum of spikes (i.e., the temporal spread of spiking) from trial-onset up to the hemodynamic response peak.

**Fig 4.**
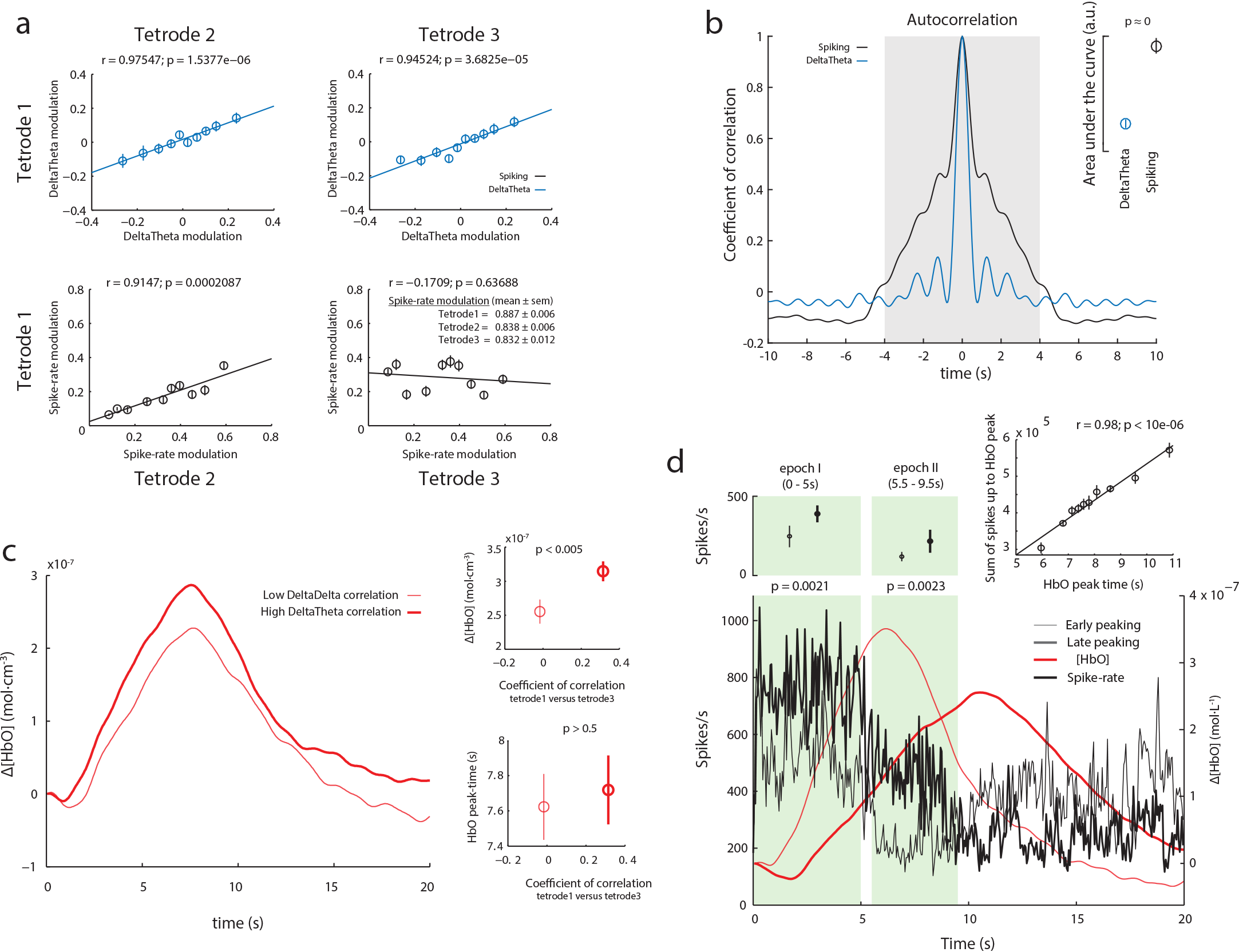
Spatio-temporal profiles of spiking versus DeltaTheta activity, and thier effect on the hemodynamic response. (a) DeltaTheta modulations are highly correlated between both Tetrodel and Tetrode2, and Tetrodel and Tetrode 3. However, spike-rate modulation are highly correlated between Tetrodel and Tetrode2, while no correlations exist between Tetrodel and Tetrode3, even though the distributions of spike-rate modulations are almost identical across tetrdodes. This illustrates that spiking activity is more spatially localized than DeltaTheta activity. (b) Mean autocorrelation of spike-rates and DeltaTheta activity across all trials. Inset) Distributions of the area-under-the-curve (between −4s to +4s, shaded region in main figure) for spiking and DeltaTheta activity. Bars represent CI-95 Spiking has a larger temporal spread than DeltaTheta. (c) Trials with higher correlations of DeltaTheta between Tetrode1 and Tetrode 3 (thick traces) have a significantly higher peak-amplitude (inset top-right) (but not peak-time (inset bottom-right) than those with low correlations (thin traces). (d) When comparing early-peaking (thin traces) vs late-peaking (thick traces) trials (top and bottom 10% of all trials, n=26), we find that late-peaking trials have significantly higher spike-rates not only during stimulus presentation (epoch I) but also after the stimulus is turned off (epoch II). A very strong correlation is observed when between the HbO peak-time and the sum of the spikes up to HbO peak-time for each trial (inset)

Curiously, we also observed that the timing of the initial-dip correlated with bursts in spike-rates better than its amplitude. To determine if the amplitude was more reflective of the spatial-spread of spiking than the strength of spiking, we analyzed the effect of synchronized bursts in spiking across tetrodes on both the initial-dip peak-amplitude and peak-time. We first obtained all trials with high spike-rate bursts on Tetrode1 (bursts larger than the median value), and further divided these trials into two groups, based on whether Tetrode2 had spike-bursts larger or smaller than the median (obtaining trials with high (n=81) and low (n=38) spike-coherence, respectively). We found that compared to low spike-coherence, trials with high spike-coherence had significantly larger initial-dip amplitudes (p<0.001; Wilcoxon’s one-tailed rank-sum test), but their initial-dip peak-times, and response peak-amplitudes and peak-times were unchanged (p>0.4) (*supplementary figure 7*). These observations support our recent suggestion that the initial-dip is a vascular response (Zaidi et al, submitted), given its dependence on the spatio-temporal dynamics of the underlying spiking activity, and also clearly demonstrate that differences in the spatial spread of neuronal processes across cortical tissue are reflected in different parameters of the hemodynamic signal.

Since we use simultaneous epidural fNIRS and intra-cortical electrophysiology to measure hemodynamic changes in anesthetized monkeys, it might be questioned how these results compare to the awake preparation, or to different neuroimaging modalities. But stark differences are unlikely, as under the anesthesia conditions we used, the correlations between neuronal activity and hemodynamic signals are not very different between the anesthetized and awake preparations, at least in the primary visual cortex (Goense and Logothetis, 2008). Furthermore, previous work in awake human subjects, where BOLD-fMRI responses to an almost identical visual stimulation paradigm were recorded, demonstrates that hemodynamic response peak-times and initial-dips are strongly correlated (Watanabe et al., 2013). Our data also reveals a strong correlation between HbO peak-time and initial-dip (r=0.84, p=0.0025; n=25/group, 10 groups), both of which correlate strongly with spiking (**Fig. 2c_1_–c_2_**, **Fig. 3b_1_–b_2_**). Based on this similarity with awake human recordings, we expect our results to be consistent across both neuroimaging modalities and states of wakefulness.

Since we do not find a relationship between excitatory neural activity and the hemodynamic response peak-amplitude, a result consistent across recordings of stimulus-induced and spontaneous activity, our study adds to the mounting evidence that the amplitude of the hemodynamic response is an unspecific marker of excitatory neuronal activity. Although we did find correlations in the hemodynamic signal’s peak-amplitude and low-frequency LFP activity, these relationships were not consistent across recordings of stimulus-induced and spontaneous activity, probably because of the difference in the underlying relationships of the various LFP bands in these two conditions (*Supplementary figure 8*). Consequently, these results also question the usefulness of canonical HRFs, as well as the practice of convolving them with neuronal activity and correlating them to observed hemodynamic responses. Furthermore, we also find that spiking is inversely correlated to low-frequency LFPs, and also to the hemodynamic response amplitude. Previous work has also demonstrated that low-frequency LFPs are inversely correlated to spiking, in both alert, behaving monkeys (Panagiotaropoulos et al., 2013) and the awake and REM states in humans (Destexhe, 1998), corroborating our results, and supporting the idea that they represent two distinct neural processes.

To summarize, our results demonstrate that the amplitude of hemodynamic responses is a poor correlate of spiking activity. Instead, we demonstrate that it is the timing of both the initial-dip and the hemodynamic response that is a much more reliable correlate of spiking activity, reflecting bursts in spike-rate and the sum-of-spikes until the hemodynamic response peak, respectively.

## Acknowledgements

The authors thank Cristina Risueno and Rebekka Bernard for help with data collection, and Vishal Kapoor and Esther Florin for feedback on an earlier version of this manuscript. This work was supported by funding from the Max Planck Society (MPG), German Research Foundation (DFG) and the Werner Reichardt Center for Integrative Neurosciences (CIN), Tübingen, the New INDIGO grant fund and the Badenwuerttemberg-Singapore Life Sciences grant.

## Abbreviations

[HbO]: : concentration of oxy-hemoglobin (mol·cm^−3^)
[HbR]: : concentration of deoxy-hemoglobin (mol·cm^−3^)
BOLD: : blood-oxygen level dependent signal
FNIRS: : functional near-infrared spectroscopy
LFP: : local field potential
MUA: : multi-unit activity
HbO peak-amplitude: : peak amplitude of the HbO response
HbO peak-time: : time from stimulus onset until peak of HbO response
SDU: : Standard-deviation units

## Methods

### Surgery and craniotomy

Two healthy adult monkeys, M1 (female; 8 kg) and M2 (male; 10 kg), were used for the experiments. All vital parameters were monitored during anaesthesia. After sedation of the animals using ketamine (15 mg/kg), anaesthesia was initiated with fentanyl (31 μg/kg), thiopental (5 mg/kg), and succinylcholine chloride (3 mg/kg), and then the animals were intubated and ventilated. A Servo Ventilator 900C (Siemens, Germany) was used for ventilation, with respiration parameters adjusted to each animal’s age and weight. Anaesthesia was maintained using remifentanil (0.2-1 μg/kg/min) and mivacurium chloride (4-7 mg/kg/h). An iso-osmotic solution (Jonosteril, Fresenius Kabi, Germany) was infused at a rate of 10 ml/kg/h. During the entire experiment, each animal’s body temperature was maintained between 38.5 °C and 39.5 °C, and SpO2 was maintained above 95%. Under anaesthesia, a craniotomy was made on the left hemisphere of the skull to access the primary visual cortex. During each experiment, the bone was removed to create a rectangular slit measuring 3 mm anterio-posteriorly and 20 mm medio-laterally, exposing the dura. Connective tissue, if present above the dura, was carefully removed. For each monkey, at least two weeks were allowed to pass between successive experiments. All experiments were approved by the local authorities (Regierungspräsidium, Tübingen) and are in agreement with guidelines of the European Community for the care of laboratory animals.

### Near-infrared Spectroscopy

We used a NIRScout machine purchased from NIRx Medizintechnik GmbH, Berlin. The system performs dual wavelength LED light-based spectroscopic measurements. The wavelengths used were 760nm and 850nm with LEDs operating at 2.5mW per wavelength. Sampling was performed at 20Hz. We used modified emitters and detectors, and optical fibre bundles for sending the light from the LED source into the tissue, and also for detecting refracted light from the tissue. The fibre bundles were ordered from NIRx Medizintechnik GmbH, Berlin, Germany. Both the emitter and detector fibre bundles had iron ferrule tips with an aperture of 2.5mm on the ends that touched the dura. The recording instrument was connected via USB to a laptop computer running an interactive software called NIRStar provided along with the instrument. The software was used for starting and stopping recordings, and also for setting up the various parameters, such as, the number of sources and detectors, and the sampling rate. The instrument received TTL pulses from the stimulus system and the electrophysiological recording system, for synchronization purposes. The system sent 1ms TTL pulses every 50ms to the recording system that corresponded to light pulses.

### Electrophysiology

Custom built tetrodes and electrodes were used. All tetrodes and single electrodes had impedance values less than 1 MΩ. The impedance of each channel was noted before loading the tetrodes on to the drive, and once again while unloading the tetrode after the experiment, to ensure that the contacts were intact throughout the duration of the experiment. To drive the electrodes into the brain a 64-channel Eckhorn matrix was used (Thomas Recording GmbH, Giessen, Germany). The electrodes were loaded in guide tubes a day before the experiment. On the day of the experiment, the tetrodes were driven using a software interface provided by Thomas Recording GmbH, Giessen, Germany. The output was connected to a speaker and an oscilloscope, with a switch to help cycle between different channels. We advanced electrodes into the cortex one by one until we heard a reliable population response to a rotating checkerboard flickering at 0.5Hz.

### Spontaneous activity

For each run, spontaneous activity was recorded for 15 minutes, in the absence of visual stimulation. The eyes of the monkey were closed and a cotton gauze was used to cover the eyelids. Peak spike-rates were obtained by implementing peak-detection in the trace of spike-rates. For spiking and all frequency bands, it was ensured that peaks are at least 20s apart from each other.

### Visual stimulation

A fundus camera was used to locate the fovea for each eye. For presenting visual stimulation, a fibre optic system (Avotec, Silent Vision, USA) was positioned in front of each eye, so as to be centered on the fovea. To adjust the plane of focus, contact lenses (hard PMMA lenses, Wöhlk, Kiel, Germany) were inserted to the monkey’s eyes. We used whole-field, rotating chequerboard to drive the neural activity. The direction of rotation was reversed every second. Each trial consisted of 5 seconds of visual stimulation followed by 15 seconds of a dark screen. A single run consisted of 20 trials. Data presented are from 14 runs spread over 8 experimental days.

### Signal processing and data analysis

All analyses were performed in MATLAB using custom written code. Only runs that cleared visual screening for artifacts were used. Data points that were larger than 5 SDU were excluded from the analysis, so as to avoid tail-effects for correlation analysis. Normality for each distribution was confirmed before analysis was performed. Trials with HbO peak-time and peak-amplitude larger than 5 SDU were removed from the analysis to avoid tail effects in correlation analysis (Supplementary figure 10).

### FNIRS signal processing

The raw wavelength absorption data from the NIRS system was converted to concentration changes of [HbO] and [HbR] using a modified Beer-Lambert equation. For correlating hemodynamic signals with neural activity, the signals were filtered between 0.01 and 5 Hz to remove any low frequency drifts. Since the change in [HbR] was negative, the peak amplitudes for [HbR] are, thus, also negative. For a trial-by-trial analysis, the hemodynamic response for each trial was zero-corrected by subtracting, from each hemodynamic response, the value at the start of the trial.

### Electrophysiological signal processing

The mean extracellular field potential signal was recorded at 20.8333 kHz and digitized using a 16-bit AD converted. From the raw signal, eight frequency bands (namely, DeltaTheta (1-8 Hz), Alpha (9-15 Hz), Spindle (15-20 Hz), low Gamma (20-40 Hz), Gamma (40-60 Hz), high Gamma (60-120 Hz), very high Gamma (120-250 Hz) and MUA (1-3 kHz)) were band-pass filtered using a 10^th^ order Butterworth filter. The envelope for each band was then obtained by taking the absolute value of the Hilbert transform of the filtered signal. The band-envelope was then converted to standard deviation units by subtracting the mean and dividing by the standard deviation of the signal. This signal was then resampled at 20 Hz, to allow comparisons with hemodynamic signals.

Spike rates were obtained by detecting peaks in the MUA signal larger than a threshold (2, 3 or 4 SDU), and by counting the threshold-crossing events in 50ms bins. Varying the detection threshold between 2, 3 or 4 SDU did not affect the results; the modulation of spike rates was significant irrespective of spike-detection threshold. Only recordings from the tetrode closest to the emitter were used.

### Calculation of modulation indices

The ‘On’ epoch for each trial was defined as the time from 0 to 5.1 seconds. The extra 0.1s were added to accommodate for the off response. The ‘Off’ epoch was defined as the time between 5.1 to 10.1 seconds. The modulation index (MI) for each trial was then calculated using the formula: neural modulation; = (BE_on_−BE_Off_) / (BE_on_+BE_Off_); where BE_On_ is the mean band envelope during the five seconds of Stim On, and BE_Off_ is the mean band envelope during the first five seconds of Stim Off for each trial.

### Statistics

All correlation coefficients represent Pearson’s correlation coefficient and corresponding significance values.

---

**Supplementary figure 1.**
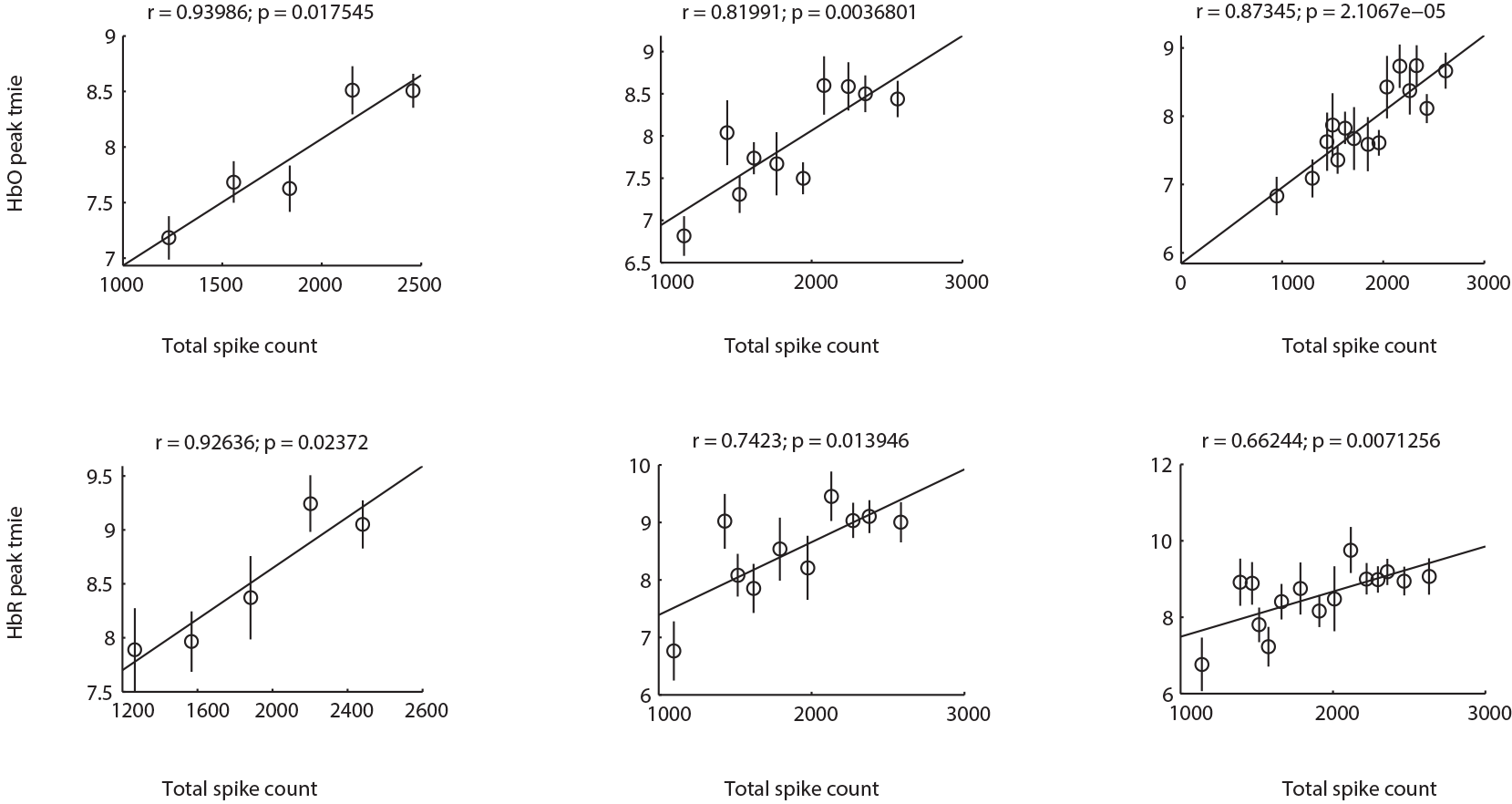
Independence of results on group size. Trials were divided into 5, 10 and 15 groups, and the mean total spike count for each group was correlated with the mean HbO and HbR peak time. Varying the size of the group did not change the relationship between the total spike counts and HbO and HbR peak times.

**Supplementary figure 2.**
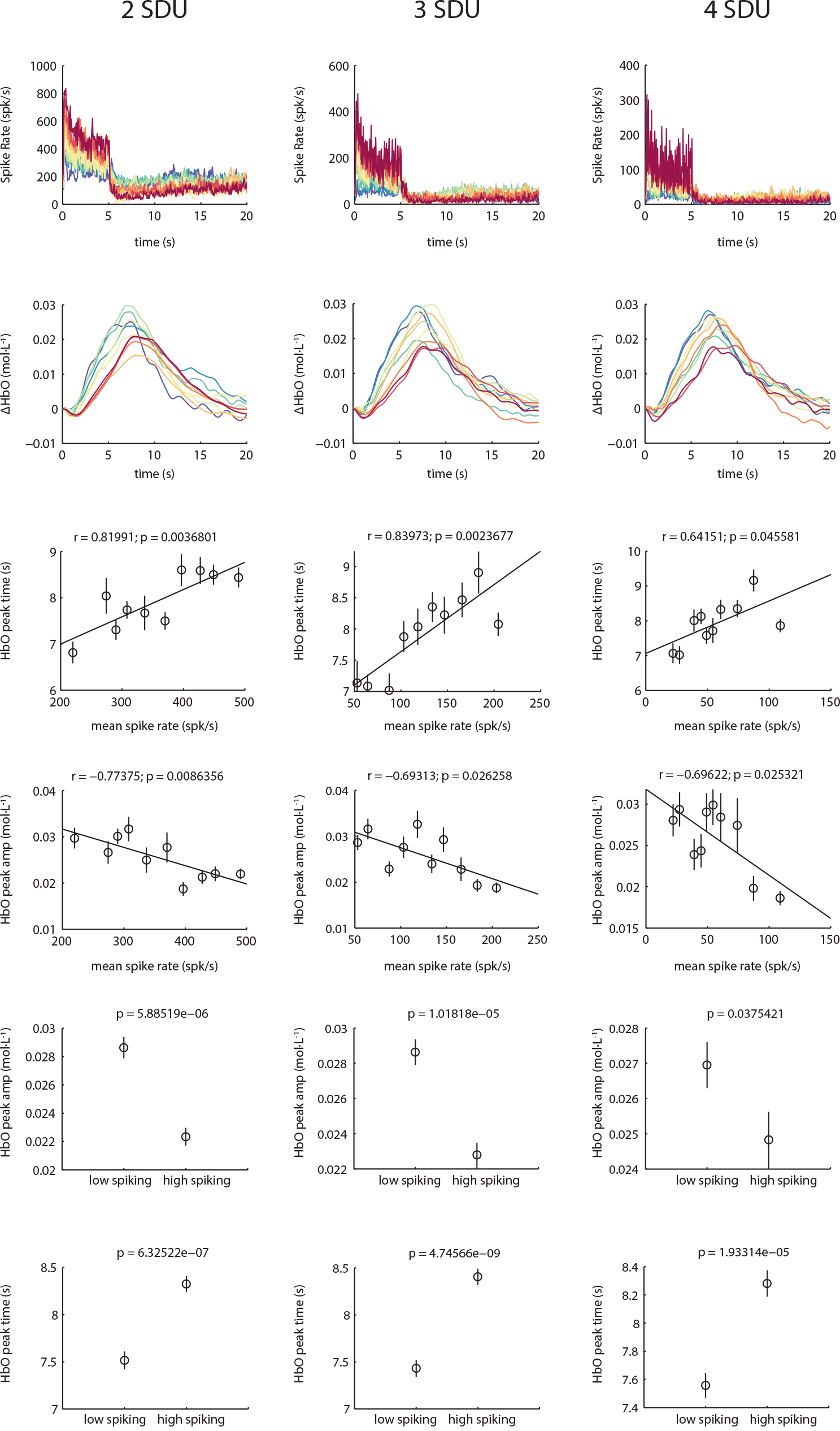
Results shown in Fig. 2 are consistent across multiple spike detection thresholds.

**Supplementary figure 3.**
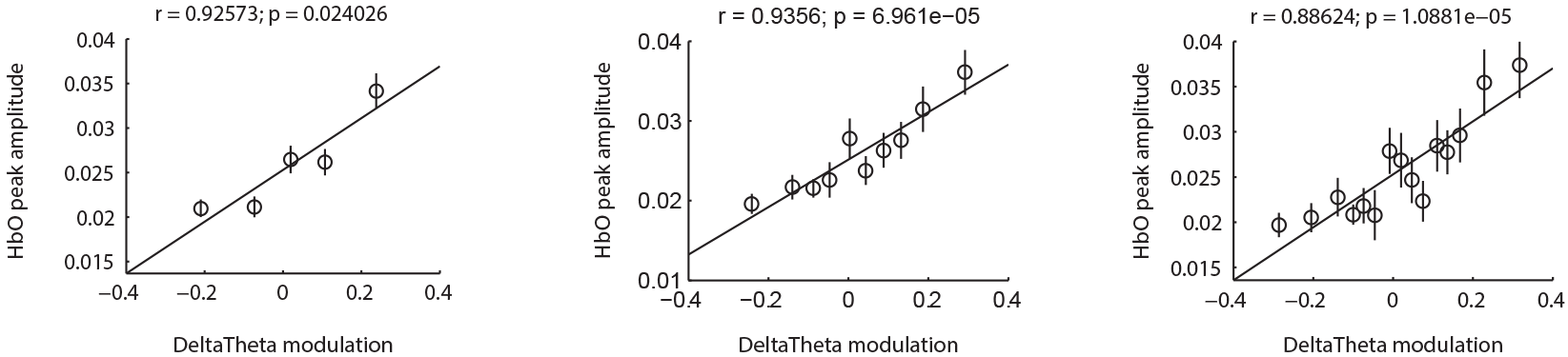
Correlations between DeltaTheta modulation and HbO peak-amplitude are independent of group size or number of trials per bin. We also observed a correlation between HbO peak-amplitude with the peak DeltaTheta amplitude during stimulus presentation (r = 0.67, p = 0.03; n=25/bin, 10 bins), but it wasn’t as strong as the correlation of HbO peak-amplitude with DeltaTheta modulation (r = 0.94, p < 10-4; n = 25/bin, 10 bins; middle panel above).

**Supplementary figure 4.**
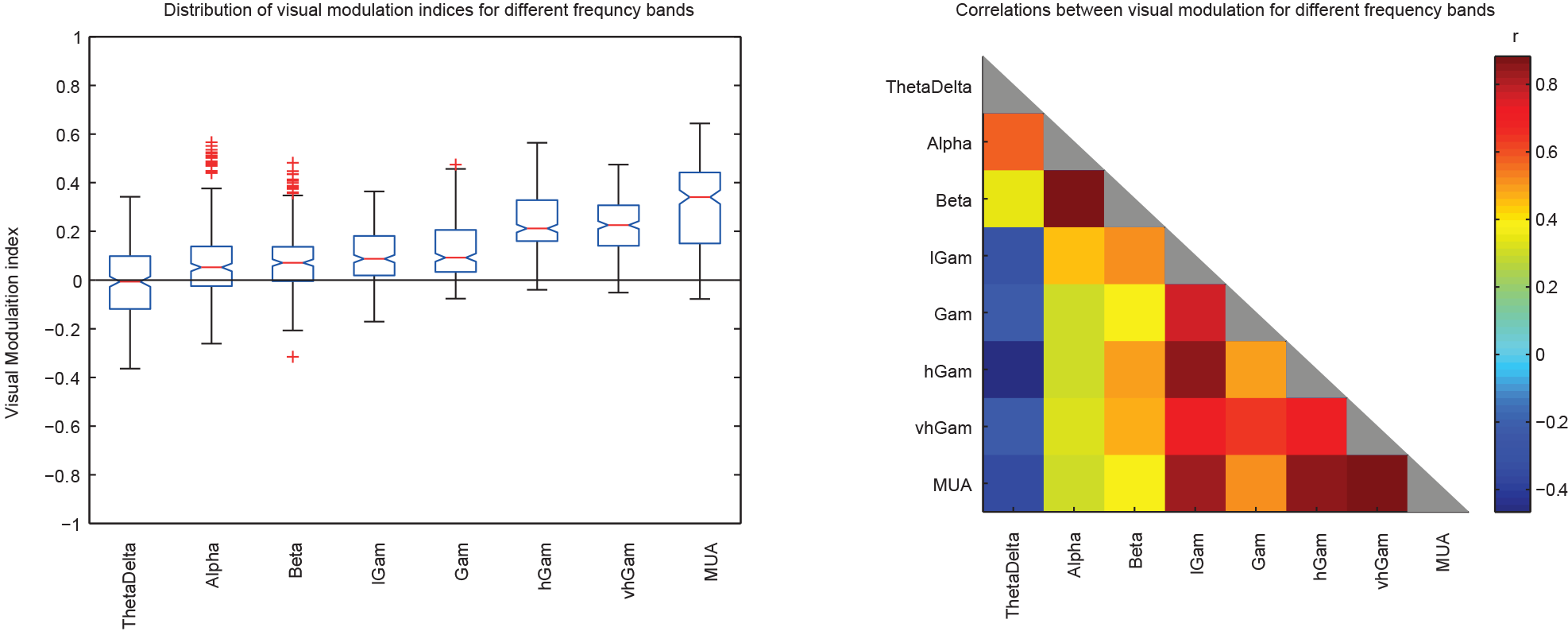
Visual modulation of different frequency bands and their relationships. Left panel shows the distribution of visual modulation indices for the different frequency bands (see Methods for definition of bands). Notches represent 95% confidence intervals. Apart from the DeltaTheta, every band had significant visual modulation, which increased with increasing frequency bands. Right panel shows the correlations between the visual modulation indices in different frequency bands. In general the three lowest frequency bands, namely DeltaTheta, Alpha and Spindle seemed to correlate positively with each other, and negatively with all bands afterwards. Modulation indices between the highest frequency bands were tightly correlated. This observation suggests that low-frequency (1-20 Hz) and high-frequency (20 - 1000 Hz) neuronal activity may represent different neuronal processes, with spiking being more representative of stimulus induced activity and low-frequency LFPs representing neuromodulatory inputs.

**Supplementary figure 5.**
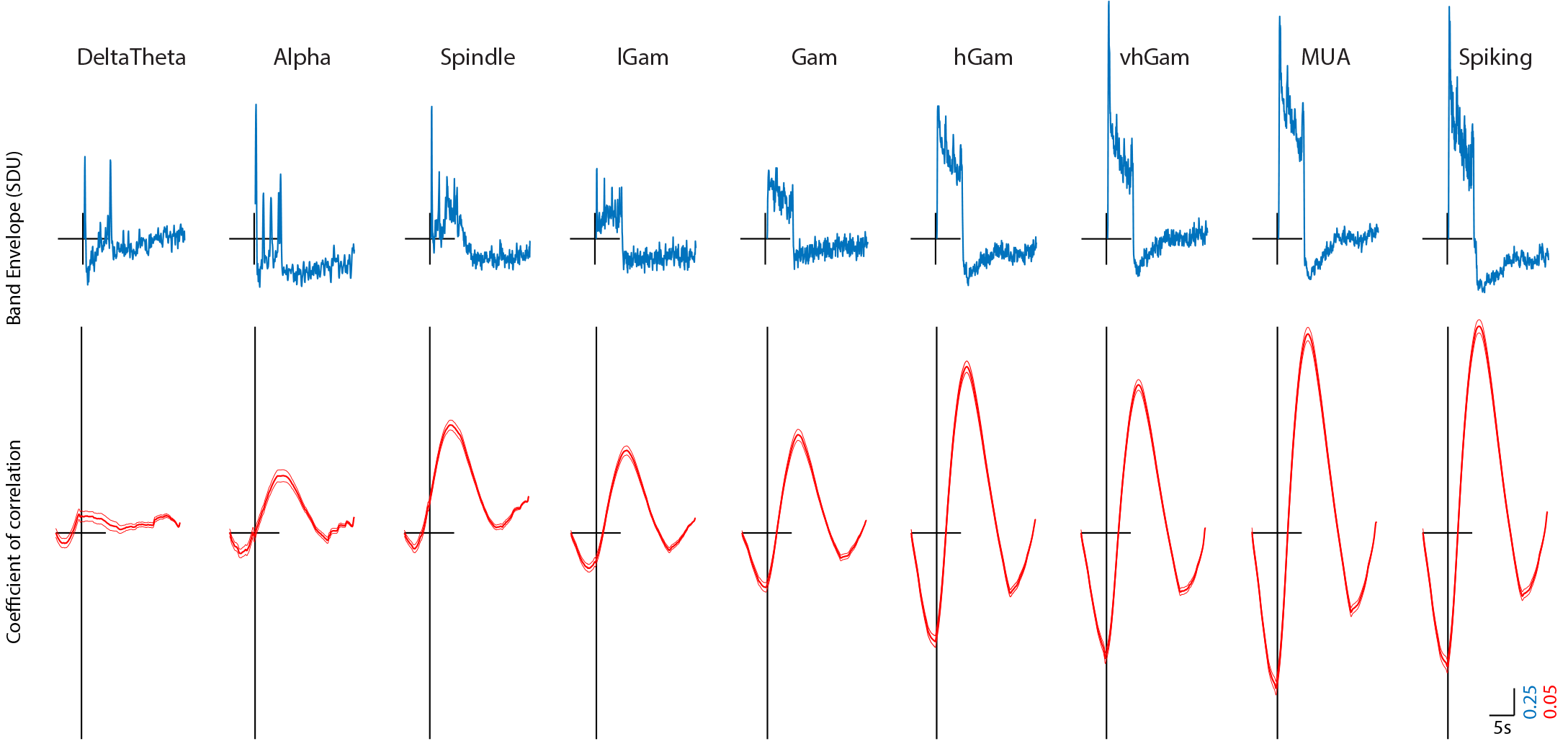
Mean LFP profiles and cross-correlations with the HbO response. The upper panels represent the mean LFP profiles for each LFP band during stimulus induced activity. The center of the cross represents the origin. The lower panels reprsent the mean cross-correlations between each LFP timeseries and the HbO response for each trial. As can be clearly observed, the cross-correlations are low for the low-frequency LFPs and high for the high-frequency LFPs. These results are fundamentally different from what we observe based on a feature-to-feature correlation analysis (see Fig. 2).

**Supplementary figure 6.**
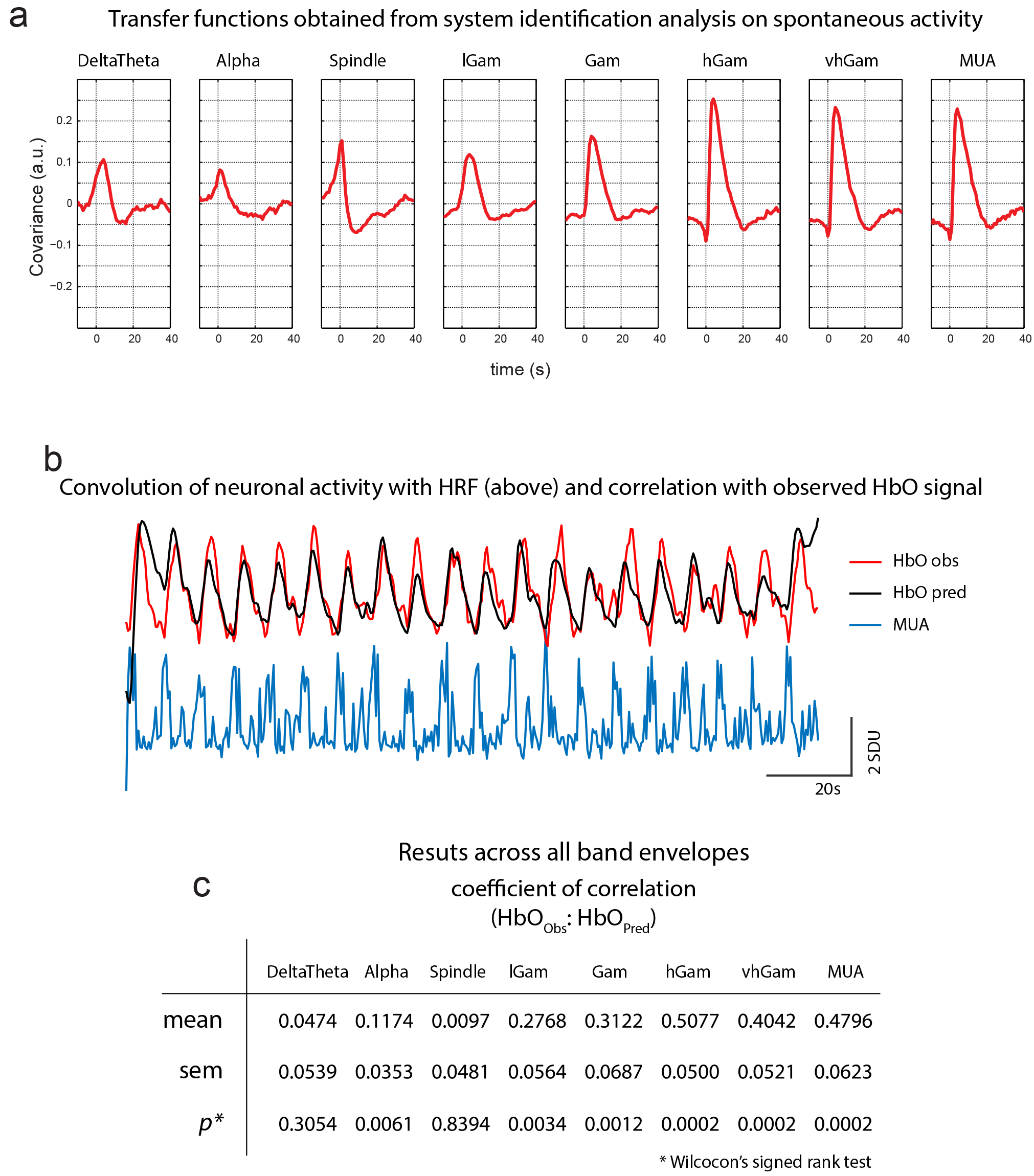
System identification based HRF estimation and prediction of stimulus induced HbO. a)The system identification toolbox in Matlab was used to obtain transfer functions (HRFs) between the band-envelope of each LFP band and the HbO signal from recordings of spontaneous activity. Each figure represents the mean HFR for that band. b) The HRFs were then convolved with each run of stimulus induced activity. c) The table represents the mean Pearson⊠ coefficient of correlation for each band⊠ along with the standard error and p-value. ⊠s can be observed⊠ the higher fre⊠uencies elicited stronger HRFs from the spontaneous activity and also stronger correlations between the predicted and observed HbO during stimulus induced activity.

**Supplementary figure 7.**
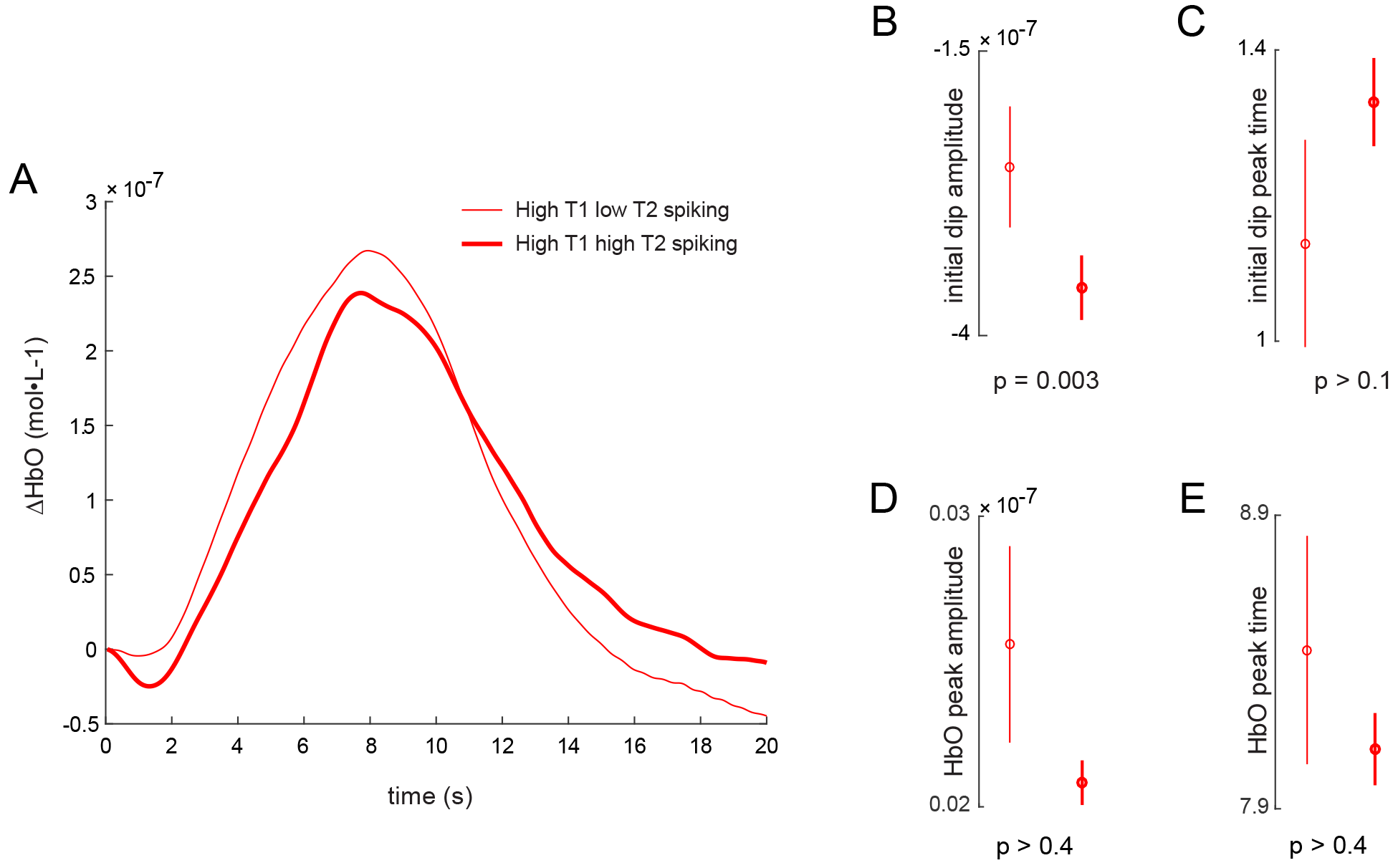
Larger spread of spike-bursts correspond to larger initial-dip amplitudes. A) The mean HbO responses for all trials with high spiking on Tetrode 1 and low (thin) and high (thick) spiking on Tetrode 2. As can be clearly observed, the initial-dip is larger for the high-spiking trials (B), without affecting either the initial-dip peak time (C), or the HbO peak amplitude or peak-time (D,E). Interestingly, larger initial dips did not correspond to larger HbO peak-times in this case, demonstrating that the initial-dip does not affect the overall HbO peak-amplitude or peak-time.

**Supplementary figure 8.**
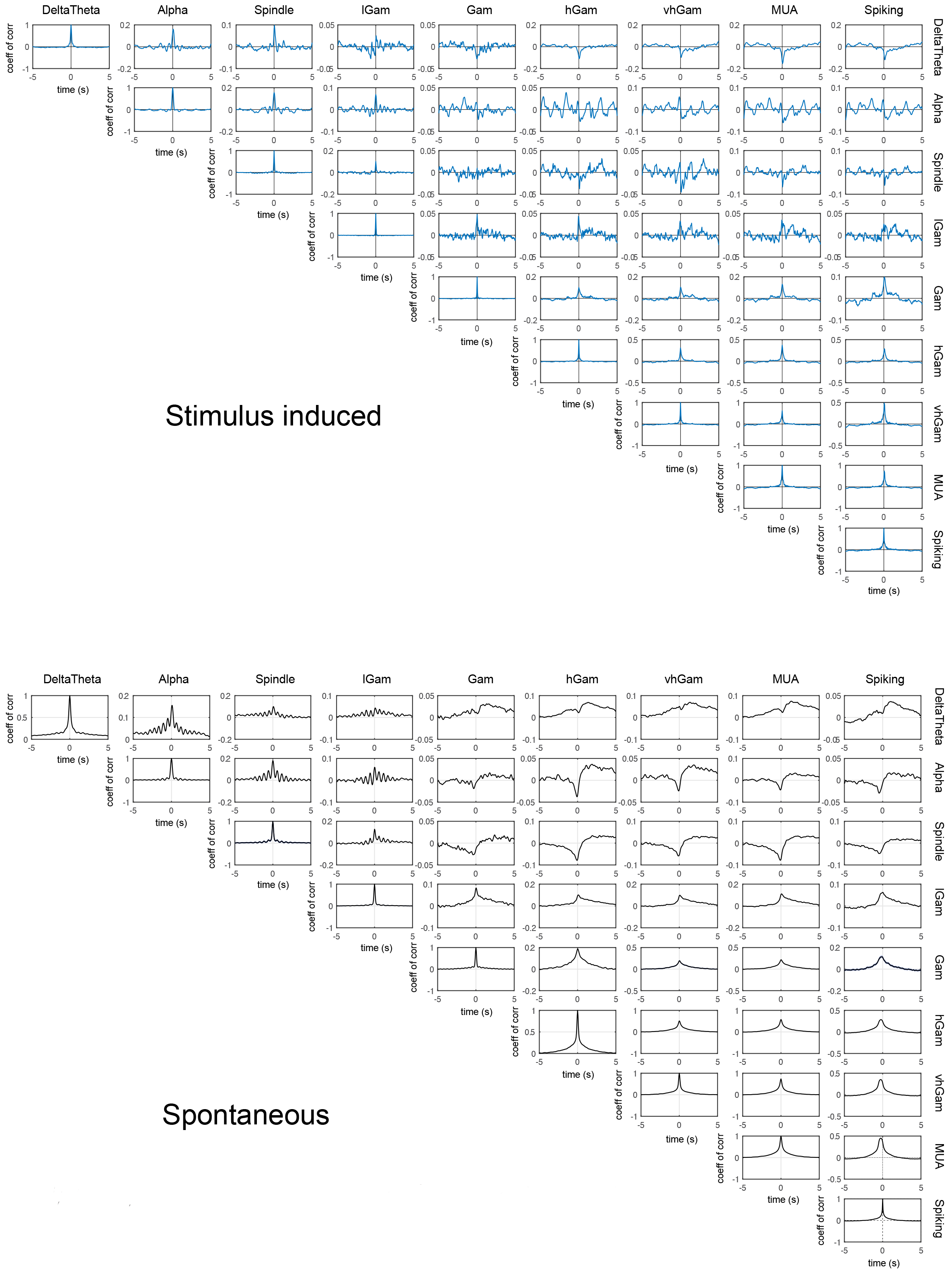
Mean cross-correlation of the band-envelopes during spontaneous and stimulus induced activity.

